# A comparison of automated and manual co-registration for magnetoencephalography

**DOI:** 10.1101/711747

**Authors:** Jon M. Houck, Eric D. Claus

**Affiliations:** Center on Alcoholism, Substance Abuse, and Addictions, University of New Mexico, Albuquerque, New Mexico, United States of America; Mind Research Network, Albuquerque, New Mexico, United States of America

## Abstract

Magnetoencephalography (MEG) is a neuroimaging technique that accurately captures the rapid (sub-millisecond) activity of neuronal populations. Interpretation of functional data from MEG relies upon registration to the participant’s anatomical MRI. The key remaining step is to transform the participant’s MRI into the MEG head coordinate space. Although both automated and manual approaches to co-registration are available, the relative accuracy of two approaches has not been systematically evaluated. The goal of the present study was to compare the accuracy of manual and automated co-registration. Resting MEG and T1-weighted MRI data were collected from 90 participants. Automated and manual co-registration were performed on the same subjects, and the inter-method reliability of the two methods assessed using the intra-class correlation. Median co-registration error for both methods was within acceptable limits. Inter-method reliability was in the “good” range for co-registration error, and the “good” to “excellent” range for translation and rotation. These results suggest that the output of the automated co-registration procedure is comparable to that achieved using manual co-registration.

## Introduction

Magnetoencephalography (MEG) is a neuroimaging technique that accurately captures the rapid (sub-millisecond) activity of neuronal populations. Indeed, MEG can only detect signal from the synchronous firing of neuronal populations in a cortical patch of approximately 10 mm^2^ or larger (1), making it essentially a network-detection technique. Due to a relative scarcity of reimbursable MEG-based clinical procedures, historically MEG was available at only a relatively limited number of cutting-edge research and clinical institutions (2). However, interest in MEG has grown as the technique’s potential has been revealed over the past four decades, with increased recognition of MEG as a means of directly evaluating neuronal networks and their relevance to a range of disorders as well as to typical cognitive and affective processes.

As is the case for other functional neuroimaging approaches, interpretation of functional data from MEG relies upon registration to an anatomical or template MRI (3). Because the data for the two modalities are collected on different scanners and therefore in different coordinate spaces (4), the procedure for MRI-MEG co-registration is somewhat involved. Typically during preparation for an MEG scan, three to five head position coils are affixed to the participant’s scalp, and then a 3D digitizing pen is used to digitize important points including the coil locations, anatomical landmarks that typically include the nasion and preauricular points, as well as a detailed headshape using approximately 150 points (Fig 1). The headshape points are collected primarily from the brow, bridge of the nose, and skull, avoiding the lower jaw and cartilaginous or fatty tissue that would be expected to shift when the participant moves or might be compressed by the head coil during the participant’s MRI scan. Because this preparation process can be somewhat labor-intensive and subject to variability in technician skill, alternatives such as use of a bite bar (5), 3-D camera (6), or 3-D laser scanner (7) have also been explored but are not widely used.

**Fig 1.**
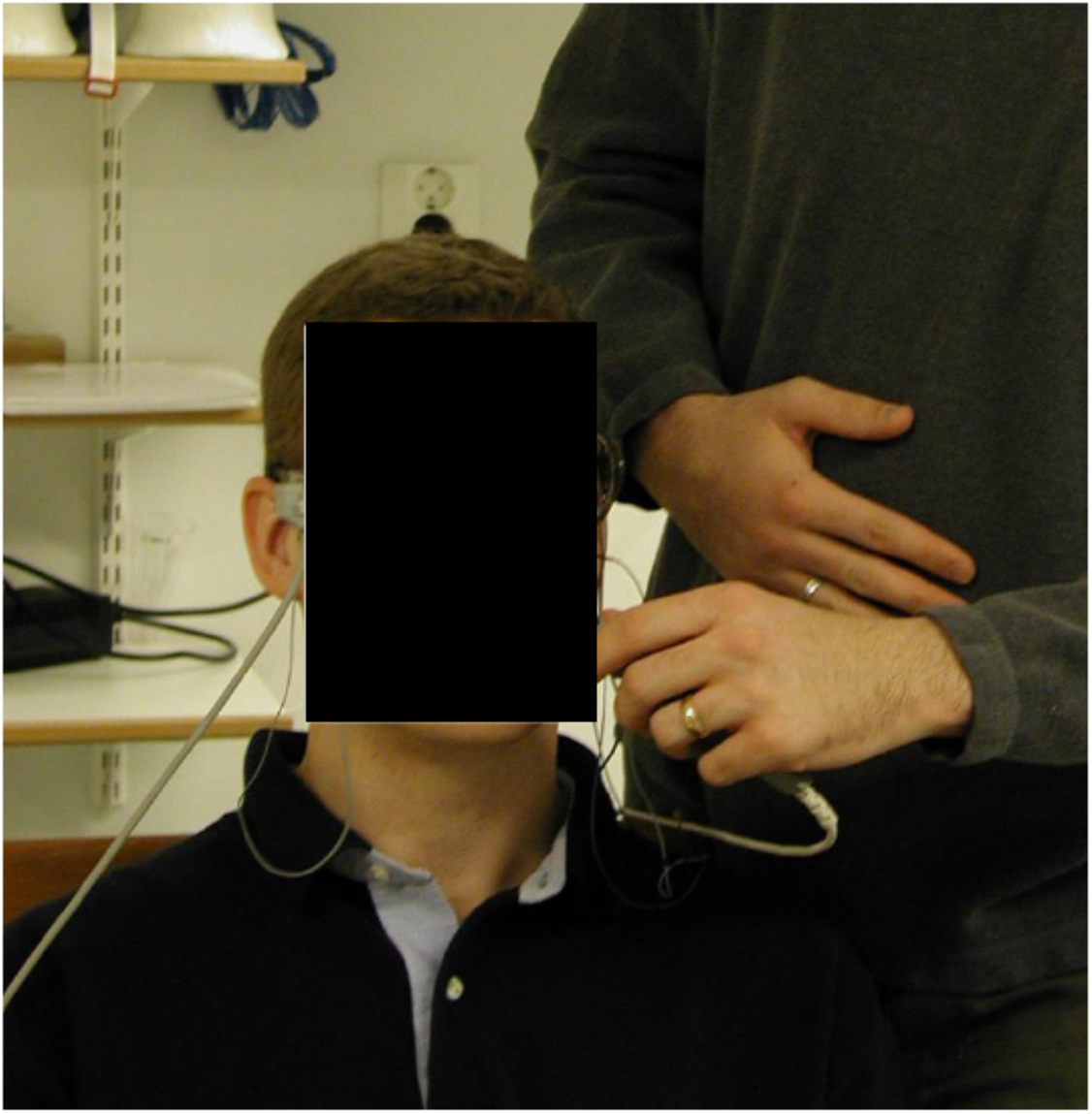
Example of collection of headshape points during participant preparation.

Subject preparation and co-registration procedures are important because they influence the quality of electromagnetic source localization (8,9). Research using MEG has consistently shown that poor co-registration quality can lead to poor source localization (10,11). When source localization is performed using beamforming, co-registration error greater than approximately 2 mm may yield unacceptably large errors in both source localization (12) and source extent (13). The same 2 mm threshold appears to apply to source localization using minimum-norm estimates (14), suggesting this threshold as an heuristic for co-registration quality.

During the MEG scan session, the head position coils are energized at known frequencies, which permits the precise measurement of their locations relative to the MEG sensor array. Because at the conclusion of an MEG scan the relative locations of the sensors, coils, anatomical landmarks, and headshape points are known, transforming the MEG data to the participant’s head coordinate space is relatively simple. The key remaining step is to transform the participant’s MRI into the MEG head coordinate space. This transformation is the focus of the MEG-MRI co-registration process.

The manual co-registration process itself is straightforward. A high-resolution 3-D head surface based on the skin-air boundary can be extracted from a T1-weighted MRI using readily-available analysis toolkits such as Freesurfer (15). Incorporation of this surface into the co-registration process has been shown to improve the quality of the co-registration (10) and is the standard in MNE-python and its predecessor, MNE (16). In the absence of significant MRI artifacts, the participant’s distinguishing features, including the face and the anatomical landmarks collected during MEG preparation, are clearly visible on this surface. The anatomical landmarks and MEG headshape can be used to co-register the MRI head surface (and therefore the MRI data) to the participant’s head coordinate space. Typically this involves manually identifying the anatomical landmarks on the MRI head surface, using these values to perform an initial transformation, and then applying an iterative closest points algorithm (ICP: 17) to assist in refining the transformation until the distance between the MEG headshape and the MRI head surface has been minimized (Fig 2). This can be accomplished using template MRIs, but MEG data can be localized with higher confidence when the individual participant’s own structural MRI is used for their co-registration. Numerous toolkits are available to assist the analysist in co-registration, including but not limited to MNE-python(18), SPM (19), Fieldtrip (20), BrainStorm (21), and NUTMEG (22).

**Fig 2.**
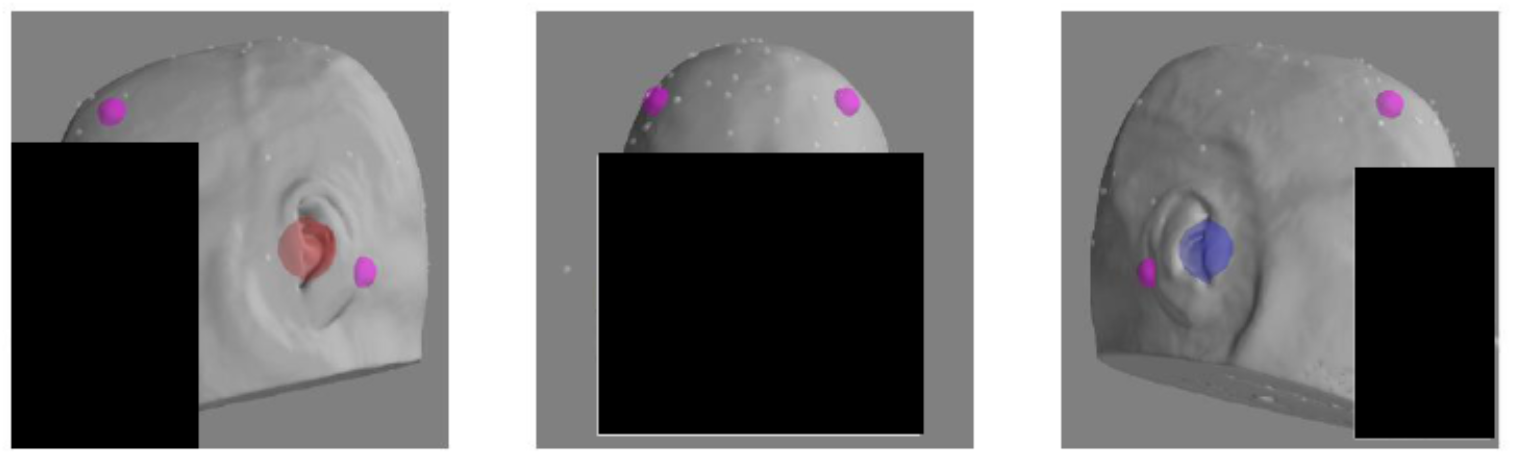
Example headshape points, landmarks, and HPI coils on head surface after co-registration.

Importantly, although both automated and manual approaches to co-registration are available, the consistency of the two approaches has not been systematically evaluated. One common approach to evaluating the consistency of two methods is to compute their inter­ method reliability. Inter-method reliability, computationally identical to inter-rater reliability, is a means of assessing the chance-corrected agreement between two different methods. This can be computed using a ratio of the variance of interest divided by the total variance; that is, the intra-class correlation (ICC: 23). The goal of the present study was to evaluate the inter-method reliability of manual and automated co-registration using the MNE-python toolbox (18).

## Method

As part of an ongoing study, resting MEG and T1-weighted MRI data were collected from 90 participants (mean age= 35.20 years (SD=10.04), 42.2% female, 50% Hispanic). MRI data were collected on a Siemens 3T Trio Tim system (Siemens Healthcare, Erlangen, Germany) using a 32-channel head coil. Paper tape was placed across each participant’s forehead to reduce motion. Structural images were collected with magnetization-prepared 180° radiofrequency pulses and rapid gradient-echo sequence (MPRAGE; TE= 1.64, 3.5, 5.36, 7.22, and 9.08 ms; TR= 2.53 s; FA= 7°; number of excitations= 1; slice thickness= 1 mm; FOV= 256 mm; resolution= 256×256). Standard preprocessing was conducted using the Freesurfer image analysis suite (15), which is documented and freely available for download online (http://surfer.nmr.mgh.harvard.edu/). However, generation of the head surface file used in co-registration relies only upon the existence of a T1 image (24), not on any specific preprocessing package.

MEG data were collected in a magnetically and electrically shielded room (VAC Series Ak3B, Vacuumschmelze GmbH) using an Elekta Neuromag whole-cortex 306-channel MEG array (Elekta Oy, Helsinki, Finland). Before positioning the participant in the MEG, four coils were affixed to the participant’s head—two on the forehead and one behind each ear. Additional positioning data was collected using a head position device (Polhemus Fastrak). Between 83 and 229 points were collected for each subject (median = 143, IQR 122-157). Participants were instructed to keep their eyes open and focused on a fixation cross back-projected onto a screen during the scan. MEG data were sampled at a rate of 1000 Hz, with a bandpass filter of 0.10 to 330 Hz. Head position was monitored continuously throughout the MEG session. Five minutes of raw single-trial data were collected and stored.

An experienced technician had previously manually co-registered each MEG scan to its corresponding MRI using MNE (16) following the general steps described in the Introduction. Automated co-registration in MNE-python follows the same general sequence described for manual co-registration. The MNE toolboxes include standard landmark coordinates (nasion, preauricular points) defined on the MNI305 head (25). The automated co-registration is performed by 1) transforming these coordinates from the MNI305 head to each participant’s MRI coordinate space, 2) performing an initial fit to the MRI head surface using only these landmarks, 3) applying several initial iterations of the iterative closest points (ICP) algorithm, 4) eliminating outlier head points (i.e., those > 5 mm away from the head surface), and 5) applying the ICP algorithm again. The final affine transformation was then saved and the co-registration errors (i.e., the median distance between each MEG headshape point and the nearest point on the MRI head surface) preserved. Errors for manual co-registrations were obtained by applying the affine transformations from the manual co-registration to the MEG headshape and computing the distance between each MEG headshape point and the nearest point on the MRI head surface. Visual inspection was used to assure the quality of the fit. The inter-method reliability of co-registration error for manual and automated co-registration was compared using the intraclass correlation (ICC model 3,1:, 23).To evaluate the relationship between participant preparation procedures and co-registration error, we computed the correlation between co-registration error terms and the number of headshape points collected during participant preparation. To assess the comparability of the head transformation matrices produced by each method, we converted each affine transformation matrix to mm of translation in the x, y, and z directions, and degrees of rotation around the x, y, and z axes (i.e., roll, pitch, and yaw), and computed the inter-method reliability for each parameter. Finally, to evaluate whether the automated technique could be applied to anonymized, “de-faced” data, we re-ran the automated co-registration procedure using MRIs that had been de-faced with PyDeface (26).

## Results

Median co-registration error for manual co-registration was 1.37 mm (IQR 1.17-1.63), and for automatic co-registration 1.58 mm (IQR 1.23-2.05) (Fig 3). The mean difference in co-registration error between manual and automated co-registration was approximately 0.313 mm (SD 0.555 mm). Co-registration error between the two methods was correlated at *r* = 0.541 (*p* < .001), which corresponds to a Cohen’s *d* of 1.29, a “large” effect size (27). The association between co-registration error and the number of headshape points was not significant for either the manual (*r* = 0.143, *p* = .058) or the automated (*r* = 0.025, *p* = 0.745) co-registration procedure. The inter-method reliability for the co-registration error between the two co-registration approaches was ICC = 0.472, which is in the “fair” range (28). After excluding automated co-registration results with unacceptably high error (i.e., > 2.0 mm), inter-method reliability improved only slightly to ICC = 0.491, also in the “fair” range (28). Inter-method reliability of all translation and rotation parameters was in the good to excellent range (i.e., all ICC < 0.74; see Table 1).

**Table 1.** Descriptive statistics and inter-method reliability results for translation and rotation *Note*. ICC = Intra-class correlation (model 3,1)

|  | Translation median in mm (IQR) |  |  | Rotation median in degrees (IQR) |  |  |
| --- | --- | --- | --- | --- | --- | --- |
|  | x | y | z | Roll | Pitch | Yaw |
| Manual | -1.861 (-4.604-0.085) | -4.842 (-9.938--<br>1.378) | -72.997 (-<br>78.808--<br>68.277) | 15.244 (9.273-19.887) | 1.394 (-0.341-3.718) | -0.955 (-2.496-0.886) |
| Automated | -1.549 (-3.543-0.514) | -5.962 (-<br>10.424--2.062) | -67.514 (-<br>71.163--<br>63.607) | 13.260 (8.011-18.450) | 1.026 (-0.668-2.687) | -0.265 (-2.135-1.276) |
| ICC | 0.903 | 0.922 | 0.833 | 0.874 | 0.748 | 0.896 |

**Fig 3.**
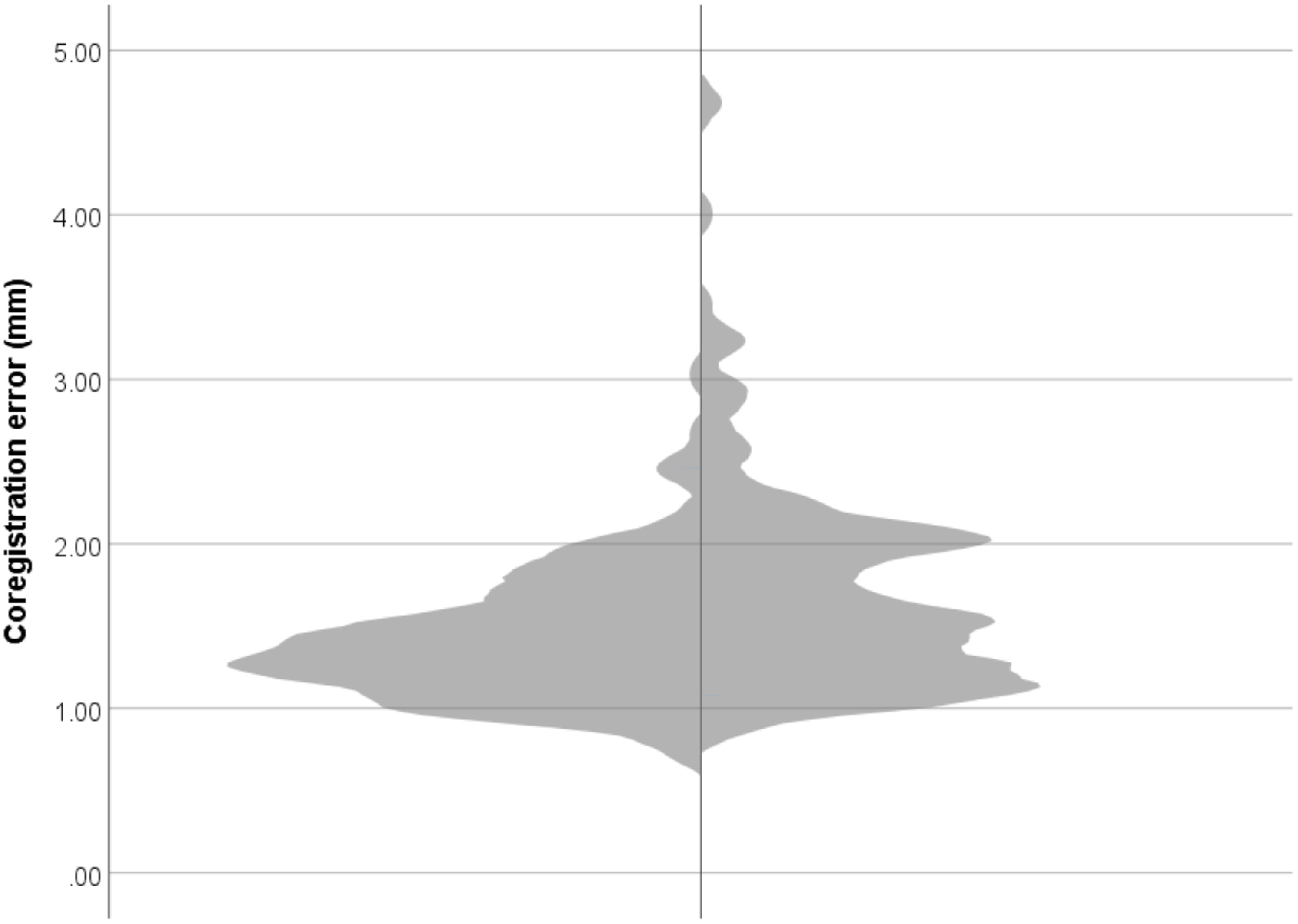
Violin plot of co-registration error for manual (left) and automated (right) approaches.

Median co-registration error for automatic co-registration with de-faced MRIs was 2.01 mm (IQR 1.71-2.32). The difference in co-registration error between manual co-registration with original MRIs and automated co-registration with de-faced MRIs was approximately 0.619 mm (SD 0.577 mm). The inter-method reliability for manual co-registration with original MRIs and automated co-registration with de-faced MRIs was ICC = 0.045, which is in the “poor” range (14).

## Discussion

In our data, both manual and automated co-registration yielded generally acceptable results. The co-registration error obtained for both processes in the present study is also consistent with that of other studies. For instance, a study using bite bars to reduce motion found a mean co-registration error of 1.16 mm (5), while a study using a 3D scanner found mean error of 2.2 mm (7), and one using a 3D camera (Kinect) observed a mean error of 1.62 mm (6). Despite the ready availability of co-registration error metrics, reporting of these metrics in MEG studies has not yet become standard practice (29,30).

The inter-method reliability results, in the “fair” range for co-registration error and the “good” to “excellent” range for translation and rotation parameters, suggests that the outputs of the manual and automated co-registration processes applied in this study are similar. That is, despite the extensive training and time requirements of manual co-registration, the results of the manual and automated co-registration procedures were in agreement, for both co-registration error and for the translations and rotations that were applied to align the MEG headshape points and MRI head surface. However, based on the results of the present study, automated co-registration using de-faced MRIs should be viewed with some caution.

It is worth noting that the MRI scans included in the present study appear to have been relatively artifact-free. Data from participants with common sources of susceptibility artifact such as braces, permanent retainers, other dental work, and certain hair products (31) would likely result in distortions of the head surface generated from the T1-weighted MRI, requiring greater attention and the potential for manual intervention during co-registration.

## Conclusion

Until devices capable of collecting simultaneous MRI and MEG data become commercially available (32), co-registration will remain a limiting factor in the localization accuracy of MEG data (10–14). Because reporting of co-registration error is not yet a best practice for MEG (29), adoption has been slow. The implementation of procedures to estimate co-registration error in analysis packages such as MNE-python (18) may help to accelerate this. Our results suggest that in many cases a simple automated processes performed using freely-available and open-source software can co-register MEG and MRI data with results similar to those achieved by manual co-registration, avoiding the time and training requirements of manual procedures.

## Acknowledgements

Research reported in this publication was supported by the National Institute on Alcohol Abuse and Alcoholism under award numbers K01AA021431 and R01AA023665. The content is solely the responsibility of the authors and does not necessarily represent the official views of the National Institutes of Health.

## Ethical approval and informed consent statement

All study protocols were approved by the University of New Mexico Institutional Review Board (http://irb.unm.edu). All procedures were carried out in accordance with the relevant guidelines and regulations. Documented informed consent was obtained from all participants. Documented consent for the publication of the photograph presented in Fig 1 and the rendering presented in Fig 2 was obtained from the individual pictured, who was not a participant in this research study.

